# Age-Related Decline in Blood-Brain Barrier Function is More Pronounced in Males than Females in Parietal and Temporal Regions

**DOI:** 10.1101/2024.01.12.575463

**Authors:** Xingfeng Shao, Qinyang Shou, Kimberly Felix, Brandon Ojogho, Xuejuan Jiang, Brian T. Gold, Megan M Herting, Eric L Goldwaser, Peter Kochunov, L. Elliot Hong, Ioannis Pappas, Meredith Braskie, Hosung Kim, Steven Cen, Kay Jann, Danny JJ Wang

## Abstract

The blood-brain barrier (BBB) plays a pivotal role in protecting the central nervous system (CNS), shielding it from potential harmful entities. A natural decline of BBB function with aging has been reported in both animal and human studies, which may contribute to cognitive decline and neurodegenerative disorders. Limited data also suggest that being female may be associated with protective effects on BBB function. Here we investigated age and sex-dependent trajectories of perfusion and BBB water exchange rate (kw) across the lifespan in 186 cognitively normal participants spanning the ages of 8 to 92 years old, using a non-invasive diffusion prepared pseudo-continuous arterial spin labeling (DP-pCASL) MRI technique. We found that the pattern of BBB kw decline with aging varies across brain regions. Moreover, results from our DP-pCASL technique revealed a remarkable decline in BBB kw beginning in the early 60s, which was more pronounced in males. In addition, we observed sex differences in parietal and temporal regions. Our findings provide in vivo results demonstrating sex differences in the decline of BBB function with aging, which may serve as a foundation for future investigations into perfusion and BBB function in neurodegenerative and other brain disorders.

**Significance statement:** The blood-brain barrier (BBB) serves as a critical protection mechanism for the CNS. A natural decline of BBB function with aging has been reported in both animal and human studies, with possible differences in BBB function by sex. Using our MRI technique, DP-pCASL that measures water exchange rate (kw) without contrast in 186 participants from diverse race and age groups, we identified age and sex-specific patterns in BBB kw especially in parietal and temporal regions. We observed of a decline in kw beginning in the early 60s, especially in males. Our study unveils the dynamic spatiotemporal pattern of kw differences with age and sex, which serve as a foundation for understanding aberrations of BBB function in neurodegenerative and other brain disorders.

## 1 Introduction

The blood-brain barrier (BBB), an intricate network of endothelial cells, pericytes, basement membrane and astrocyte end-feet, forms a selective permeability shield between the central nervous system (CNS) and blood. This dynamic interface regulates the passage of macromolecules to allow the passage of nutrients and preventing the entry of potential neurotoxins and pathogens (1). There is a natural decline and breakdown of BBB function with aging as revealed by both animal and human studies (2, 3). When combined with a second hit such as inflammation and/or ischemia, this trend of declining BBB function with healthy aging may become more detrimental leading to neuronal damage and pathogenesis of a variety of neurological disorders including multiple sclerosis (4, 5), stroke (6-8), traumatic brain injury (9), Parkinson’s Disease (10), cerebral small vessel disease (cSVD) (8, 11) and Alzheimer’s disease (12-14). Considering the prominent difference in the prevalence of these disorders between males and females, it is reasonable to speculate that there may be sex differences in BBB integrity which may vary with age. A few animal studies have reported a protective effect of female sex hormones on BBB permeability, with ovariectomized rats showing increased Evan’s blue dye extravasation into the brain which was normalized by estrogen replacement (15, 16). Two human studies showed that compared to males, females have significantly lower CSF/serum albumin ratio and reduced BBB permeability to Gadolinium (Gd)-based contrast agent using dynamic contrast enhanced (DCE) MRI (17, 18). However, lumber puncture for CSF sampling and venous injection of Gd-based contrast agent are required to measure BBB permeability in existing human studies. However, these methods have drawbacks as the CSF/serum albumin ratio does not provide regional information and DCE MRI is not suitable for specific populations (e.g. children and participants with renal dysfunction)

Recently, a non-invasive MRI technique termed diffusion prepared pseudo-continuous arterial spin labeling (DP-pCASL) (19, 20) has been proposed for mapping BBB water exchange rate (kw) by differentiating labeled water in the cerebral capillaries and brain tissue with appropriate diffusion weighting. Compared to Gd-based contrast agents, water is an endogenous tracer with a small molecular weight (18 Da), and water exchange across the BBB occurs at a relatively high level and is mediated by passive diffusion, active co-transport through endothelial membrane, and facilitated diffusion through the dedicated water channel, aquaporin-4 (AQP4), at the end-feet of astrocytes (21, 22). As a result, assessing kw, which may reflect the function of AQP4 as suggested by preclinical studies (22), is expected to provide a more sensitive metric of BBB function compared to DCE MRI.

DP-pCASL has recently been applied in a range of brain disorders and has been validated by mannitol administration to open the BBB and using an ischemia-reperfusion model to disrupt BBB in rats (23, 24). As opposed to increased BBB leakage detected by DCE MRI (25), reduced BBB kw has been found in a wide range of CNS disorders including obstructive sleep apnea (26), schizophrenia (27), multiple sclerosis (4) and heredity cSVD (28, 29). In particular, emerging evidence suggests a high relevance between BBB water exchange and glymphatic function. Amyloid plaques may interfere with the normal function of AQP-4 and lead to its depolarization (30, 31). Recent studies reported that BBB kw was lower in those with more *APOE* ε4 alleles and more amyloid deposition in the brain as detected by PET (32). Similarly, in cognitively normal adults, BBB kw was lower in those with lower CSF Aβ-42 (typically associated with the more pathological condition) (33). These data suggest associations between reduced BBB water exchange, compromised glymphatic function and impaired clearance of Aβ (34), potentially due to dysfunctional AQP4 (31, 35). Reduced BBB kw with aging has also been reported in small groups of young and aged persons (36-38), suggesting that the glymphatic system becomes less effective in older adults (39).

Despite the promising findings, the dynamic spatiotemporal pattern of BBB kw evolution with age and sex is not yet understood. The aim of the present study was to investigate potentially region-specific trajectory of kw variations with age and sex in a cohort of 186 cognitively normal participants aged from 8 to 92 years. We recruited only cognitively normal individuals to discern the influence of natural aging on BBB function, which may serve as a future foundation for detecting BBB aberrations in neurodegenerative and other brain disorders. Based on the literature and our recent findings, we hypothesized that that age-related BBB kw declines may be more pronounced in males than females.

## 2 Results

### Population

This study included 186 participants aged 8 to 92 years (89 males and 97 females) with a diverse racial distribution including 65 White/Caucasian (29 males, 36 females), 27 Latinx (11 males, 16 females), 47 African American (21 males, 26 females), and 47 Asian (28 males, 19 females) (Supplemental Fig. S1). Since race may influence the changes in perfusion and kw with aging, it was included as a covariate in the following statistical analyses.

### Age and sex-dependent trajectories of regional kw, ATT, and CBF changes

The DP-pCASL technique provided concurrent measurements of arterial transit time (ATT), cerebral blood flow (CBF), and kw (Fig. 1A) with a single scan. Anticipating that associations of regional CBF, ATT, kw with age may be nonlinear, especially if BBB degradation initiates within a specific age range, we employed Multivariate Adaptive Regression Splines (MARS) (40) analysis, which is adept at automatically identifying nonlinearities (splines) and interactions among variables with a 10-fold generalized cross validation (GCV) to prevent over-fitting.

**Figure 1.**
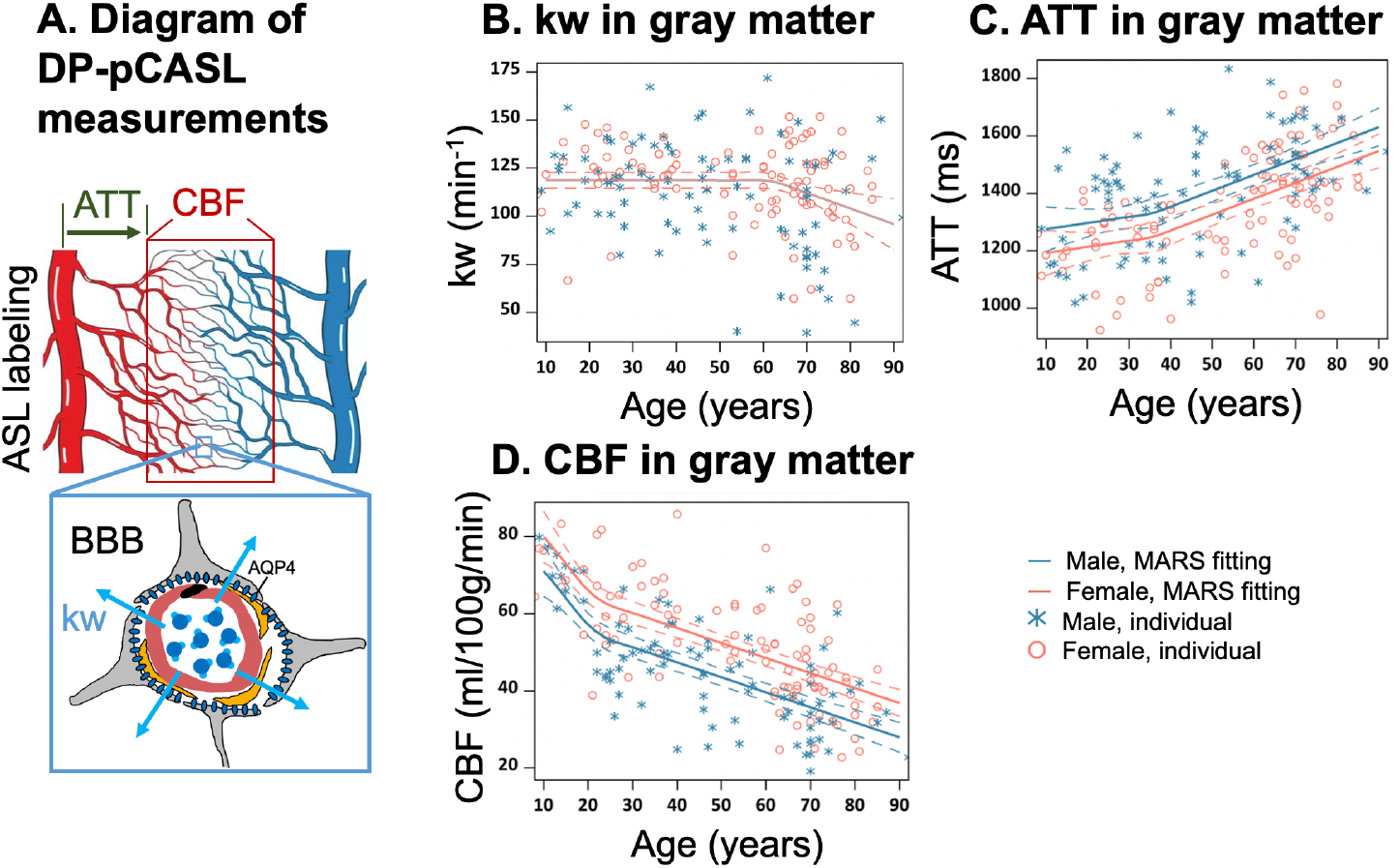
Illustration of DP-pCASL measurements and age-related trends in kw, ATT, and CBF in gray matter. (A) Diagram illustrating the DP-pCASL measurements. ATT represents the transit time of the labeled blood traveling from DP-pCASL labeling plane to the imaging voxel, CBF refers to the amount of labeled blood supplied to the brain per unit of time (or perfusion), and kw describe the rate of blood water exchanging from intravascular space (capillaries) into extravascular space (tissue) facilitated by multiple transport mechanisms including AQP-4 water channel assisted transport. (B-D) Scatter plots representing age-related distribution of kw (B), ATT (C) and CBF (D) values in the gray matter. In all scatter plots, individual data points for males and females are indicated by blue asterisk symbols and red circles, respectively, while the corresponding MARS fitting curves and 95% confidence interval for expected value at each age point are presented as continuous lines and dashed lines in matching colors.

Within the gray matter, we observed that BBB kw remained relatively consistent before the age of 62 years and declined thereafter with a yearly slope of −0.82 (95% CI: [-1.35, −0.28]) (P=0.003), while no statistically significant sex differences were observed by MARS analysis (Fig.1 B, Supplemental Table S1). In contrast, ATT was overall longer in males, while CBF was higher in females across the lifespan (Fig.1 C, D, Supplemental Table S2, S3). It is worth noting that the patterns of change in these three physiological parameters were different, with kw showing a turning point around the age of 62 years (Fig.1 B, Supplemental Table S1), ATT around the age of 36 years (Fig.1 C, Supplemental Table S3), and CBF around the age of 22 years (Fig.1 D, Supplemental Table S2). Regional MARS analysis results for all 14 subregions can be found in Supplemental Figures S3-S5 and Tables S1-S3 for kw, ATT and CBF, respectively.

Sex difference became evident for kw in MARS analyses of brain regions such as the parietal lobe (Fig. 2 A, P=0.01), temporal lobe (Fig. 2 B, P=0.001), medial temporal lobe (MTL) (Fig. 2 C, P=0.008), hippocampus (Fig. 2 D, P=0.02) and parahippocampal gyrus (PHG) (Fig. 2 E, P=0.006) (Supplemental Table S1). In these regions, kw decline with age was more pronounced in males than females. In contrast, the rates of CBF decrease and ATT increase with aging were largely consistent between the males and females, except for a more rapid CBF decrease in males within the hippocampus (Fig. 2 F, P<0.001, Supplemental Table S2).

**Figure 2.**
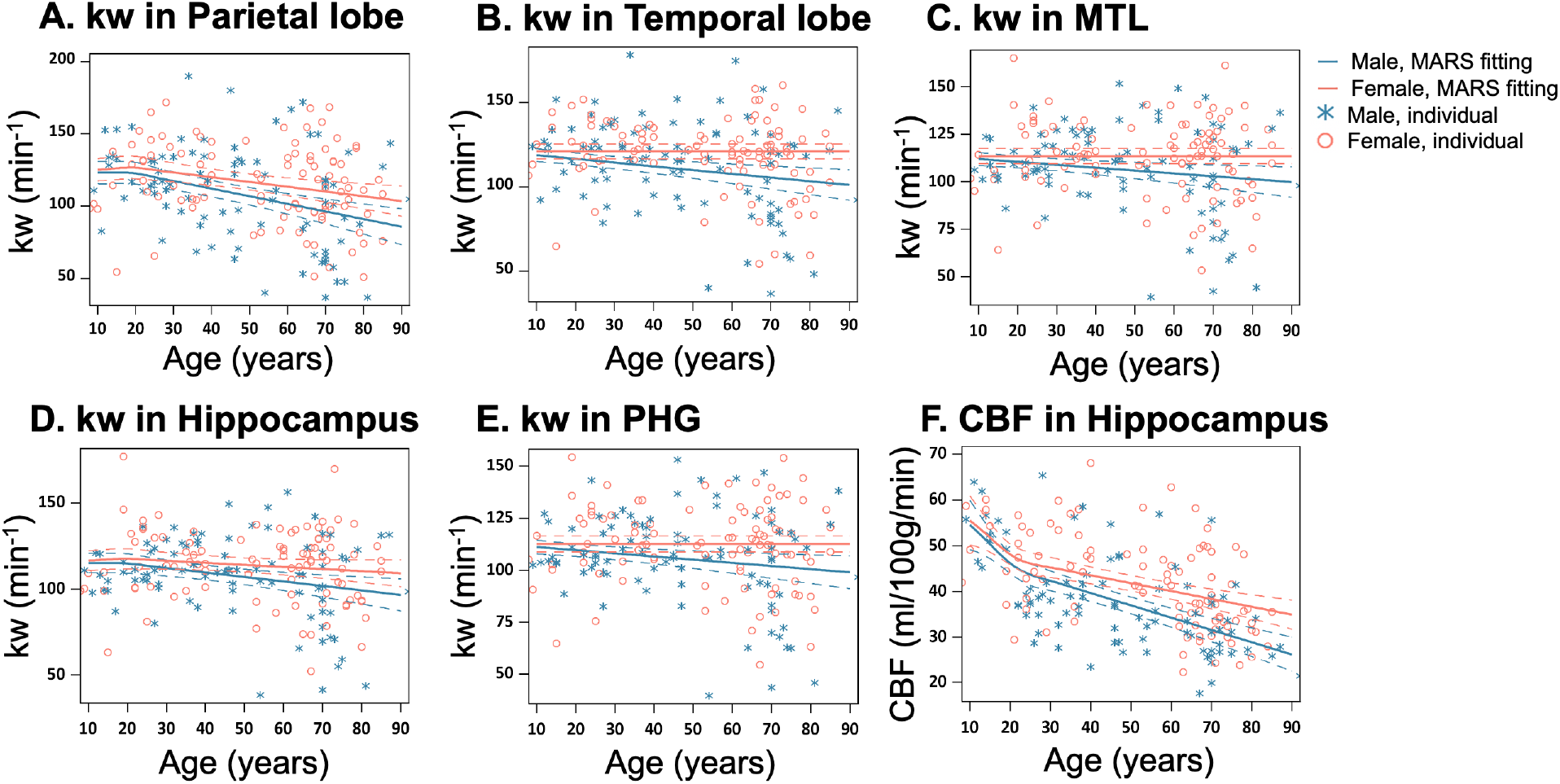
Sex specific age trends in kw and CBF. (A-E) Scatter plots representing age-related distribution of kw values in the parietal lobe (A), temporal lobe (B), MTL (C), hippocampus (D) and PHG (E). F. Scatter plots representing age-related distribution of CBF values in the hippocampus. In all scatter plots, individual data points for males and females are indicated by blue asterisk symbols and red circles, respectively, while the corresponding MARS fitting curves and 95% confidence interval for expected value at each age point are presented as continuous lines and dashed lines in matching colors. Abbreviation: MTL, medial temporal lobe; PHG, parahippocampal gyrus.

### Average kw, ATT and CBF maps across age groups

We further divided the cohort of 186 participants into 3 age groups with similar number of participants spanning a comparable age range, and calculated average maps of kw, CBF and ATT for the three age groups: children to young adulthood (8-35 years, N=56), middle age (36-61 years, N=55), and older age (62-92 years, N=75) (41) (Fig. 3). The threshold of 62 years as the starting point for the older age group was consistent with the MARS results, which indicated a notable decline in BBB kw beginning after the age of 62 years. Whole-brain average kw values for males were 120.7±17.4, 117.5±26.5, and 97.9±31.4 min^-1^ across the 3 age groups, while females’ kw values were 121.7±18.2, 118.9±14.0, and 114.8±23.7 min^-1^, respectively. Whole-brain average CBF values were 51.5±11.9, 43.9±10.7, and 31.9±8.5 ml/100g/min for males, and 60.5±10.7, 51.2±11.7, and 39.6±10.3 ml/100g/min for females from young to middle age to elderly groups. Whole-brain average ATT values were 1325.8±157.7, 1383.9±199.6, and 1526.7±117.4 ms for males, and 1220.2±134.2, 1334.4±155.4, and 1468.1±166.9 ms for females across the three age groups. These average maps of 3 age groups are highly consistent with the regional MARS analysis.

**Figure 3.**
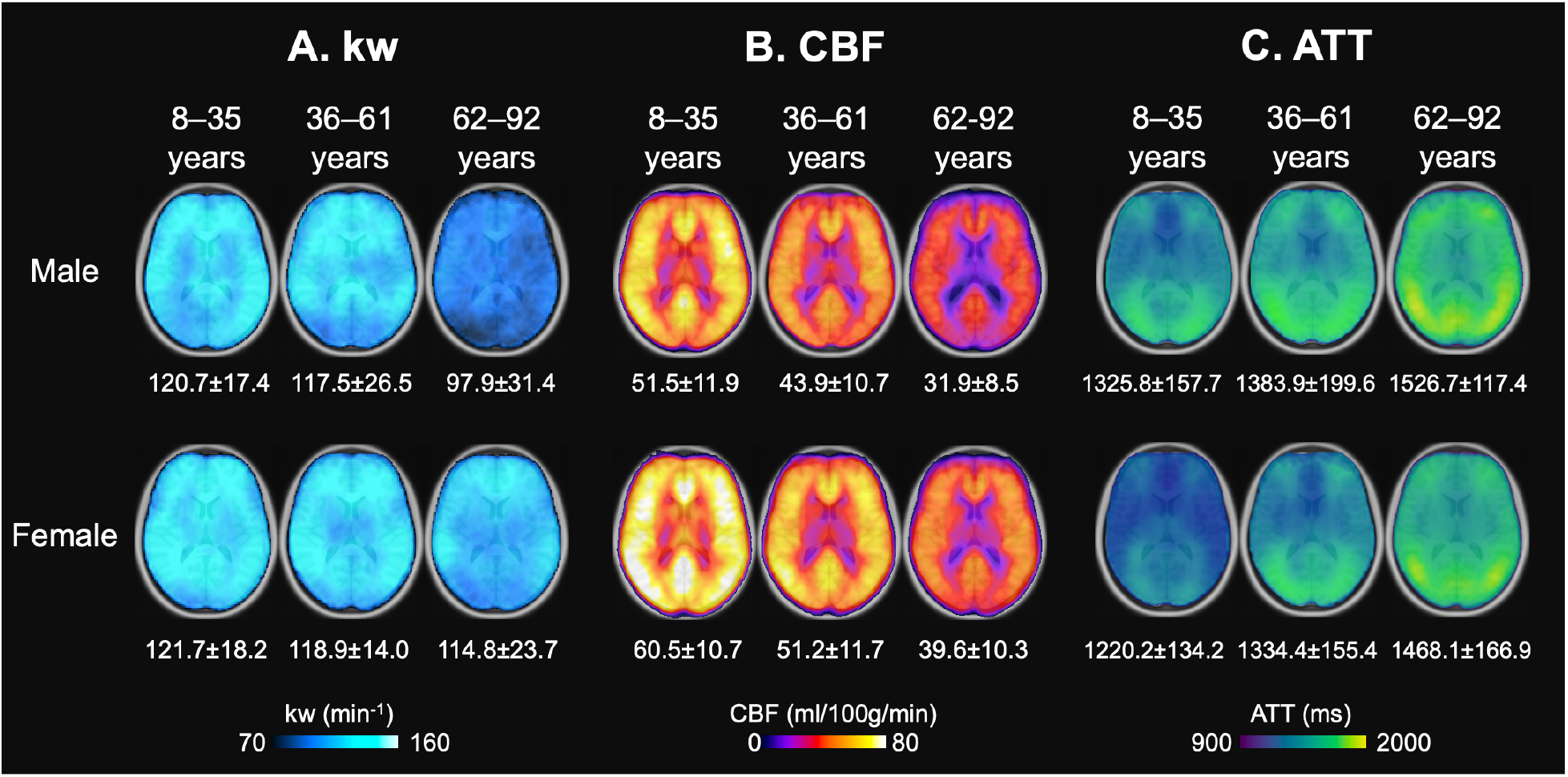
Age and Sex-based Variations in kw, ATT, and CBF. (A) kw maps, (B) CBF maps, and (C) ATT maps, average across three age groups: 8-35 years (Males: n=31, average age 23.0 years; Females: n=26, average age 22.7 years), 36-63 years (Males: n=28, average age 46.8 years; Females: n=28, average age 51.3 years), and 62-92 years (Males: n=30, average age 72.7 years; Females: n=43, average age 72.8 years). These maps are superimposed on T1w anatomical images. Corresponding average kw, CBF, and ATT values are provided beneath each map. Across the spectrum, kw values remained relatively consistent between males and females, though a marked reduction in kw can be observed in males aged 62-92 years. Patterns in the maps suggest an age-related a crease in CBF and increase in ATT, while males generally had lower CBF and longer ATT values compared to females.

As age progresses, we observed increase in ATT and a concurrent decrease in CBF, while males had 8.6%, 3.7% and 3.9% longer ATT and 14.8%, 14.3% and 19.4% lower CBF as compared to females across 3 age groups respectively (P<0.001, one-way ANOVA). The sex difference in kw values was more pronounced among older participants (62-92 years), while males had an average of 14.7% lower kw compared to females (P<0.001, one-way ANOVA) (42). Minimal differences (<2%) in mean kw values were found in young and middle-aged cohorts. Supplemental Fig. S2 A shows that in the older participants, males had slightly longer ATT (3.9%, P<0.001) compared to females. However, sex differences can be observed in kw distributions (Supplemental Fig. S2 B), where females show higher mean kw values while males present a broader distribution. To determine whether the lower kw observed in males may be attributable to prolonged ATT, we conducted simulations recalculating females’ kw values to match males’ ATT by adding 60 ms, leading to marginally higher kw values in females (Supplemental Fig. S2 C). These findings suggest that the kw differences observed between males and females are not predominantly driven by ATT differences. Furthermore, considering our SPA model’s kw quantification algorithm, which utilizes the ratio of perfusion signals with and without diffusion weightings (20), it is unlikely that CBF variations contribute significantly to the sex differences in kw. To account for potential confounding effects, we included both ATT and CBF as covariates in the following regression analyses of kw with age and sex.

### Voxel-level analysis of age and sex effects on kw, CBF, and ATT

To investigate detailed spatiotemporal pattern of perfusion and kw variations with age and sex, we further applied voxel-wise analysis using two generalized linear regression models (GLM) to study the effect of age and the interaction between age and sex on CBF, ATT, and kw respectively (see details in Materials and Methods). In particular, for voxel-wise analysis of each of the 3 parameters (e.g. kw), the rest 2 parameters (e.g. CBF and ATT) were included as covariates in the GLM to control for the potential confounding effects of the 3 parameters concurrently measured by DP-pCASL.

### Age trend

Figure 4 shows T-statistic maps of effects (GLM1), which suggested that age-related differences in CBF and ATT were widespread, encompassing the majority of brain regions (Fig. 4 B and C). Conversely, significant decreases in kw were primarily observed in specific brain regions (Fig. 4 A) including lateral and ventro-medial prefrontal cortex, cingulate cortex (CC), precuneus, medial and lateral temporal lobe, occipital lobe, and insula (Supplemental Table S4). In the brain regions showing significant age-related kw decreases (Fig. 4A), these decreases are mostly accompanied by CBF decreases (Fig. 4B) and ATT increases (Fig. 4C).

**Figure 4:**
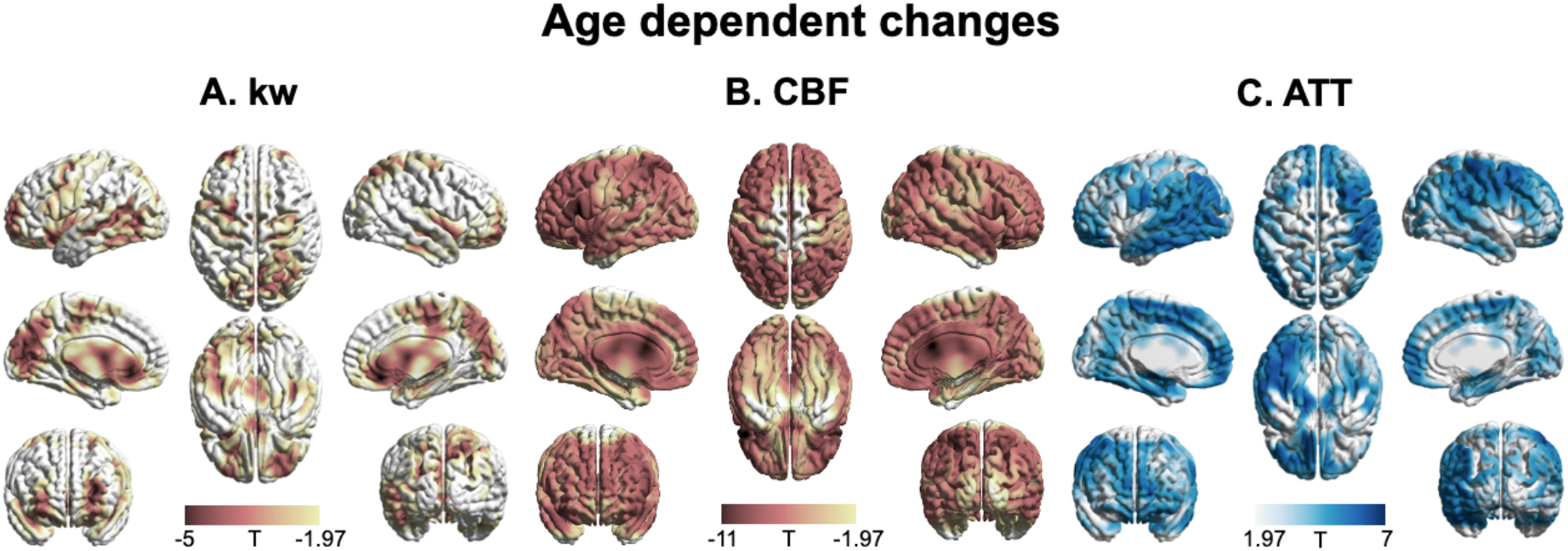
3D renderings of T maps depicting age-related differences. Highlighted areas indicate significant age-related decrease in kw (A), decreased CBF (B), and increased ATT (C). The most pronounced decrease in kw was observed in the lateral and medial prefrontal cortices, ACC, PCC, temporal lobe, parietal lobe, occipital lobe and insula. A broad range of brain regions exhibits both CBF reduction and ATT increase. The color scale represents T values, and clusters consisting of over 501 voxels with an absolute T value greater than 1.97 are considered significant and displayed in the figure. The limited slice coverage of DP-pCASL (96 mm) may account for the absence of detected effects in the upper and lower regions of the brain. The effects of decrease or increase were represented by warm colors (yellow to red) and cold (gray to blue) colors, respectively.

### Sex effect

Figure 5 displays T-statistic maps showing the interaction effects between age and sex on kw (A), CBF (B), and ATT (C), respectively. We found an accelerated decline in kw as males age in comparison to females in regions including lateral prefrontal cortex, parietal as well as lateral and medial temporal lobes (Supplemental Table S5). Although significant age dependent changes of CBF and ATT were found in majority of brain regions, very few clusters remained significant when examining the interaction effects between age and sex. We found pronounced decline in CBF with advancing age in males relative to females in specific clusters located around the supramarginal gyrus, hippocampus, and frontal lobe (Supplemental Table S5). An accelerated increase in ATT with aging in males was found in a few clusters within regions including supramarginal gyrus, posterior temporal lobe and calcarine sulcus (Supplemental Table S5).

**Figure 5.**
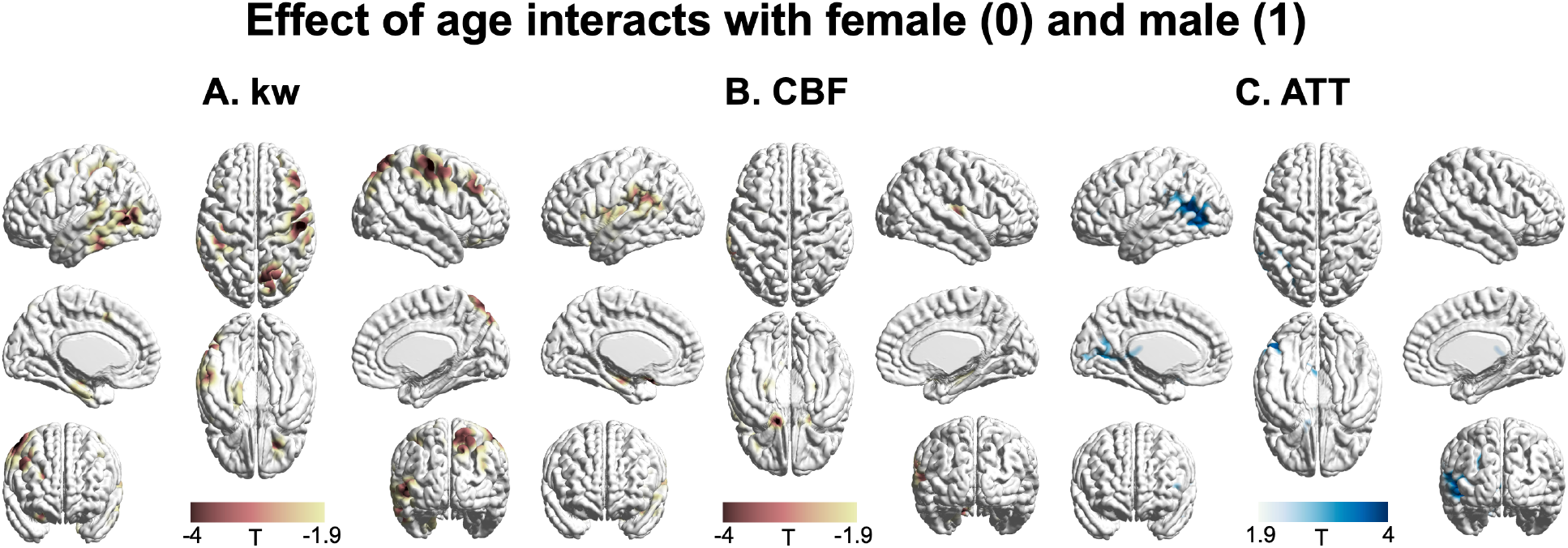
3D renderings of T maps illustrating the interaction effects of age with sex. Highlighted areas indicate: A. Accelerated decrease in kw with aging in males compared to females, most evident in the lateral prefrontal cortex, parietal, and lateral and medial temporal areas. B. Accelerated decrease in CBF with aging in males compared to females, prominently observed in the supramarginal gyrus, hippocampus, and frontal areas. C. Accelerated increase in ATT with aging in males compared to females, with marked changes in supramarginal gyrus, posterior temporal lobe and calcarine sulcus. The distinct interaction patterns between age and sex across kw, CBF, and ATT can be observed. The color scale denotes T values. Clusters comprising over 501 voxels and possessing an absolute T value exceeding 1.90 are deemed significant and showcased in the figure. The effects of decrease or increase were represented by warm colors (yellow to red) and cold (gray to blue) colors, respectively.

## 3 Discussions

To the best of our knowledge, the present study is the first to simultaneously investigate the spatiotemporal trajectories of CBF, ATT and BBB kw variations with age and sex in a diverse group of cognitively normal participants across the lifespan (27, 33), enabled by the innovative DP-pCASL technique. Our findings indicate a pervasive increase in ATT and decrease in CBF across brain regions (Fig. 4), with similar rates of change between males and females in majority of brain regions (Fig. 2 and 5). Females show higher CBF and shorter ATT compared to males of similar age, which is consistent with literature findings on CBF and ATT variations with age and sex (43, 44).

Longer ATT suggests a delayed delivery of blood to brain tissue, possibly due to multiple factors such as compromised vascular elasticity or cerebrovascular reactivity (45). Females have demonstrated relatively higher CBF values compared to males, potentially due to the protective effects of estrogen on the vasculature as well as lower hematocrit levels in females (46). Furthermore, we found a more pronounced age-related decline in CBF in the hippocampus of males compared to females (Fig. 2, Supplemental Table S2). To the best of our knowledge, no study has previously reported this accelerated hippocampal CBF decline in males. This finding may be linked to the accelerated hippocampal volume loss in males, as reported in a study analyzing 19,793 generally healthy UK Biobank participants (47). Lower hippocampal perfusion has been associated with poor memory performance (48, 49), suggesting that males might be more vulnerable to potential cognitive decline (50).

We observed a distinct trajectory of kw changes with aging compared to CBF and ATT. To study the potential regional associations between kw and CBF and ATT, we conducted linear regressions between regional kw and regional CBF or ATT, incorporating sex as a covariate, for participants aged 8-61 years and 62-92 years (when BBB kw starts declining), respectively. The results are shown in Supplemental Table S6. BBB kw was significantly negatively associated with CBF in the putamen, amygdala, hippocampus, PHG, and MTL in participants aged 8-61 years (when kw was relatively consistent across ages), but no significant correlations were found in any brain regions in the 62-92 years group. In contrast to CBF, kw was significantly negatively associated with ATT in the GM, temporal lobe, and precuneus in participants aged 8-61 years, and these correlations became significant in additional brain regions, including WM, frontal lobe, ACC, caudate, putamen, amygdala, hippocampus, PHG, and MTL in participants aged 62-92 years. These results suggest that BBB function may be affected by different aspects of neurovascular function represented by CBF and ATT at different stages of aging.

### BBB kw during normal aging

The kw remained relatively stable throughout early to mid-adulthood, with a marked decline in the early 60s, especially in males (∼18%). This pattern is consistent with previous research on age-related changes in BBB function. For instance, Sweeney and colleagues found evidence for BBB breakdown in the hippocampus, a region crucial for learning and memory, even before clear signs of cognitive dysfunction were evident (51). Similarly, Montagne and colleagues associated early BBB dysfunction with later cognitive impairment, highlighting the vulnerability of the aging brain to vascular changes (52). Finally, Taheri and colleagues reported that AQP4 function can differ with age and might contribute to changes in kw (53).

We found that specific brain regions are more susceptible to BBB kw decline with aging including regions within each of the four lobes as well as regions that span multiple lobes such as the insula, and cingulate gyrus. BBB dysfunction in this widespread set of these regions could negatively affect a broad range of cognitive processes including executive functions (54), memory processing (55, 56), emotional regulation (57, 58), sensory perception (59, 60). Our findings with cognitively normal participants suggest that BBB dysfunction, as indicated by a decline in the kw metric, might precede detectable cognitive impairment. Our results thus suggest that non-invasive imaging techniques such as DP-pCASL may make a significant contribution to future clinical trials seeking to identify and enroll individuals at-risk for vascular cognitive impairment for emerging pharmacological treatments.

There is a natural decline and breakdown of BBB function with aging as revealed by both animal and human studies (2, 3), which affects all BBB components (endothelial cells, astrocytes, pericytes, microglia, neuronal elements). Other studies reported that the BBB function is relatively stable during early to mid-adulthood but may show alterations with aging (13). Several studies have reported that BBB dysfunction and leakage starts to increase in middle-aged individuals and becomes more pronounced in the elderly, especially in the presence of neurodegenerative diseases (1, 52, 61, 62). However, the exact age at which BBB dysfunction begins is still under debate and may vary between individuals. Age-related changes in BBB function can be due to a combination of factors, including reduced expression of tight junction proteins (63, 64), increased endothelial cell leakage and permeability (65), decreased efflux transporter activity (66), and inflammatory changes (67). An emerging role of AQP4 on BBB function and glymphatic system has been discovered recently (68), and expression and polarization of AQP4 channels can change with aging, which can affect water homeostasis and potentially contribute to age-associated neuropathologies (69-71). Our finding of a general trend of kw decline with aging beginning in the early 60s is consistent with the molecular mechanisms of BBB decline and breakdown during healthy aging.

### Decline in BBB function is more pronounced in elderly males

The most intriguing finding of our study is that BBB kw declines faster in males compared to females in specific brain regions including lateral prefrontal and parietal cortex, as well as lateral and medial temporal lobes. The lateral prefrontal cortex and parietal cortex are heavily inter-connected and together support executive functions, planning, and reasoning (72). The medial temporal lobe plays a vital role in memory encoding and retrieval (73). We also observed an asymmetric effect on left and right brain hemispheres, which might be associated with asymmetrically developed vascular burdens in aging (74). Our findings align with evidence indicating that women might maintain certain types of memory skills better than men as they age, suggesting a sex-based resilience in areas of the brain associated with cognitive functions (75, 76).

Sex differences in susceptibility to BBB dysfunction, may be influenced by multiple factors including hormonal, genetic and lifestyle factors (77). A protective effect of female sex hormones on BBB permeability has been reported, with ovariectomized rats showing increased Evan’s blue dye extravasation into the brain which was normalized by estrogen replacement (15, 16). However, the benefit of estrogen replacement has been observed only in young but not in aged rats (78). In humans, females show lower CSF/serum albumin ratio and reduced BBB permeability to Gd-based contrast agent using DCE MRI compared to males (17, 18). Estrogen is known to have a neuroprotective effect through several mechanisms (79). The distribution of estrogen receptors on the surface of endothelial cells varies across brain regions, contributing to the observed region-specific kw trajectories between males and females. Additionally, age-related decline in estrogen levels or alterations in estrogen receptors may contribute to the disruption of BBB integrity (80).

Another possibility is that the sex effect we see is related to genetics. Recent studies underscore the X chromosome’s integral role in immune function, suggesting its gene expressions may contribute to the sex differences observed in BBB function (81, 82). Genes, such as tissue inhibitor of matrix metalloproteinase (TIMP), which occasionally escape X-chromosome inactivation, have been linked to protect BBB disruption (83). Elucidating the genetic underpinnings may illuminate the pathogenesis of BBB-related dysfunctions, with implications for understanding sex differences in neurological disorders.

Additionally, lifestyle factors may contribute to the observed sex differences in BBB integrity. For instance, the prevalence of sleep apnea, which increases with age in men, has been associated with compromised glymphatic clearance (84). This could partially explain the lower BBB kw observed in older males.

Moreover, understanding the interplay between sex and BBB function might have broader implications. For instance, while estrogen is known to confer some neuroprotective effects (85), this might not be the sole factor responsible for observed sex differences in BBB dysfunction, especially in older age (86, 87). While the precise mechanisms behind these sex differences in BBB evolution with age remain a topic of ongoing investigation (88), our study and the DP-pCASL technique offer a completely noninvasive approach to probe the mechanisms underlying these age and sex-related changes in kw, potentially paving the way for early detection and intervention for cognitive decline and neurodegenerative disorders.

### Limitations of the study and future directions

There are a few limitations of this study. A single PLD of 1800 ms was used in this study, which should be sufficient to allow all the labeled water to reach the tissue (i.e., the longest ATT was 1526.7±117.4 and 1468.1±166.9 ms in aged males and females, respectively) (19). However, a longer PLD should be used in participants with longer expected ATT, such as in stroke and cerebrovascular disorders. Additionally, a multi-PLD protocol can also be helpful to improve the robustness of quantification accuracy (89). To compensate for the half signal loss of the non-CPMG DP module, relatively low spatial resolution and TGV-regularized SPA modeling were employed. Our recently development of a motion-compensated diffusion weighted (MCDW)-pCASL can be utilized to improve the spatial resolution in the future studies (e.g. 3.5 mm^3^ isotropic maps in 10 mins) (89). Mahroo et al., utilized a multi-echo ASL technique to measure BBB permeability to water and reported shorter intra-voxel transit time and lower BBB exchange time (Tex) in the older participants (≥50 years) compared to the younger group (≤20 years) (90). In animal studies, reduced BBB Tex was also reported in the older mice compared to the younger group using multi-echo ASL (91) and a multi-flip-angle, multi-echo dynamic contrast-enhanced (MFAME-DCE) MRI method (92). These findings contrast with the results presented in this study, likely due to the different components assessed by different techniques, and increased BBB permeability to water has been suggested to indicate a leakage of tight junctions in aging (91, 92). In contrast, our recent study utilizing high resolution MCDW-pCASL scans with long averages reveals the potential existence of an intermediate stage of water exchange between vascular and tissue compartments (e.g., paravascular space or basal lamina) (89). The DP module of the DP-pCASL is hypothesized to null the fast-flowing and pseudo-random oriented spins, which may include both vascular flow and less restricted water in paravascular space. The observed lower kw in older participants may be more related to the delayed exchange across the astrocyte end-feet into the tissue due to loss of AQP-4 water channel with older age. However, these hypotheses require further investigation to understand the exact mechanisms, especially under different physiological stages (38, 93). Future studies, particularly with animal models targeting specific BBB components under different physiological or diseased conditions, will be valuable for validating these measurements (6, 22, 94-96). Including race as a covariate in our study aims to account for potential variations in brain perfusion observed in previous research (97, 98). However, it is important to recognize that these differences may not be solely attributable to race. They can be influenced by a complex interplay of factors such as education, environmental exposures, lifestyle, healthcare access, and other social determinants of health (99). For example, education has been shown to be highly relevant to regional CBF changes in AD (100, 101). Additionally, the potential influence of ancestry and mixed-race on perfusion and BBB function requires further investigation in future studies. Other factors such as hematocrit (23), menopausal status (24, 25), and vascular risk factors (26) should also be considered. These variables were not included in this study due to the unavailability or limited availability in some cohorts. We attempted to minimize the impact of these factors on our observations by including a relatively large and diverse sample. However, future studies examining the specific mechanism of each of these factors on BBB function in aging would be valuable.

### In conclusion

simultaneous mapping of perfusion and BBB kw using DP-pCASL revealed pronounced declines in BBB kw across the lifespan especially in males in their early 60s, as well as sex related difference in specific brain regions. Our findings on age and sex related trajectories of perfusion and BBB variations may provide a foundation for future investigations into perfusion, BBB function and glymphatic system in neurodegenerative and other brain disorders.

## 4 Materials and Methods

### MRI measurement of BBB kw using DP-pCASL

All participants were scanned on 3T Siemens Prisma scanners using 32 or 64 channel head coils. The imaging parameters of DP-pCASL were: resolution = 3.5 × 3.5 × 8 mm^3^, TR = 4.2s, TE = 36.2 ms, FOV = 224 mm, 12 slices+10% oversampling, labeling duration = 1.5 s, b=0 and 14s/mm^2^ for post-labeling delay (PLD) of 0.9s with 15 measurements, b=0 and 50 s/mm^2^ for PLD of 1.8s with 20 measurements, and total scan time = 10 mins. In addition, 3D MPRAGE for T1 weighted structural MRI (TR=1.6s, TI=0.95s, TE=3ms, 1mm^3^ isotropic spatial resolution, 6 min, parameters vary slightly between participating sites) were acquired for segmentation and coregistration.

### Study cohorts

We included 207 cognitively normal participants without current or past major medical illnesses, head injury with cognitive sequelae, intellectual disability, or current substance abuse (MOCA≥26 and CDR=0 for elderly participants, middle aged and young participants were screened for any neurologic or psychiatric disorders or intellectual disability). Participants were recruited from the following cohorts including University of Southern California (USC) MarkVCID study (https://markvcid.partners.org/) for elderly Latinx participants, USC Sickle cell trait (SCT) SCT study for African American participants, University of Maryland, University of Kentucky and USC for racially diverse participants including young to middle-aged to elderly participants. Among recruited participants, 21 of them were excluded due to lack of demographic information or severe image artifacts. A total of 186 participants (97 females and 89 males) with age ranging from 8 to 92 years old were included for further analysis, including 65 white, 27 Latinx, 47 African American and 47 Asian participants. Detailed distributions of age, sex and race are shown in Supplemental Fig. S1. All participants were asked to avoid caffeine at least 3 hours ahead of the scan and stay awake during the DP-pCASL scan. Additional head padding was used to reduce head movement during the scan. For more vulnerable participants (participants under 17 years old), we had instructed them to stare at the cross shown on the screen behind the scanner to minimize head motion.

### Image preprocessing

DP-pCASL data was analyzed using our in-house-developed LOFT BBB toolbox including motion correction, physiological noise reduction using principle component analysis (PCA) (102), and calculation of signal differences between the labeled and control images. ATT maps were generated using DP-pCASL signals acquired at the PLD = 0.9 s with b=0, 14 s/mm^2^ (FEAST method) (103) and CBF calculated using the PLD = 1.8 s data according to the single-compartment Buxton model (104). The kw map was generated with total-generalized-variation (TGV)-regularized single-pass approximation (SPA) model using DP-pCASL signals acquired at the PLD = 1.8s with b=0, 50 s/mm^2^, as well as CBF, ATT and tissue relaxation time (R_1b_) map generated from background suppressed control images (20). Intracranial volume (ICV) and voxel-wise gray matter (GM) density was obtained by segmentation of T1w MPRAGE images using SPM12 (http://fil.ion.ucl.ac.uk/spm/). CBF, ATT and kw maps were co-registered to the T1w MPRAGE images along with M0 image of DP-pCASL, and then normalized into the Montreal Neurological Institute (MNI) standard brain template (2 mm^3^ isotropic resolution) using SPM12 before group-level analysis.

### Regional analysis using MARS

Anticipating that associations of regional CBF, ATT, kw with age may be nonlinear, especially if BBB degradation initiates within a specific age range, we employed MARS (40) for analysis, which is adept at automatically identifying nonlinearities (splines) and interactions among variables. To mitigate the risk of overfitting, we conducted 10-fold Generalized Cross Validation (GCV), aiming to minimize outlier influence and reduce model complexity. This approach ensures an optimal prediction model boasting the best generalizability. The final MARS model, utilizing the hockey stick basis function, yielded predicted values for the testing sample, with each prediction grounded in models trained on mutually exclusive data sets following the 10-fold cross-validation. The final model performance was presented as the R^2^ between model predicted and observed value. The predicted spline slope with corresponding 95% confidence interval for expected value at each age point was illustrated using scatter plot with spline trajectory curve. We have fit the MARS model for each imaging feature (CBF, ATT, kw) in 14 subregions including GM, WM, frontal lobe, temporal lobe, parietal lobe, anterior cingulate cortex (ACC), posterior cingulate cortex (PCC), precuneus, caudate, amygdala, hippocampus, parahippocampal gyrus (PHG) and medial temporal lobe (MTL) based on the Automated Anatomical Atlas (AAL) template (105). Unlike the conventional statistical modeling, MARS is a machine learning tool with the output mainly focuses on the predicted accuracy from cross-validation. To facilitating the model interpretation, the basis functions selected by MARS as the important predictors were refitted in generalized linear model to obtain the slope estimation with 95% confidence interval. SAS 9.4 was used for MARS model fitting and validation.

### Voxel-wise analysis using GLM

We employed two GLM models to study the voxel-wise effect of age and the interaction effects between age and sex on CBF, ATT, and kw. The models are detailed as follows:

### Age effect analysis (GLM1)

This model was designed to investigate the influence of age on the dependent variables. The equation for GLM1 is as follows:

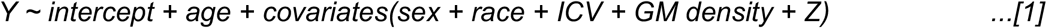

### Interaction effect analysis of age and sex (GLM2)

GLM2 aimed to analyze the interaction effects between age and sex on the dependent variables. The equation for GLM2 is expressed as:

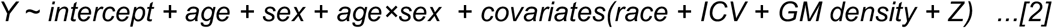

In both models, ‘Y’ represents an individual imaging feature (either CBF, ATT, or kw), while the remaining two features are included as covariates (represented as ‘Z’ in equations [1, 2]). For instance, when analyzing kw changes, ‘Y’ denotes kw, and ‘Z’ includes CBF and ATT.

For the statistical analysis, we corrected for multiple comparisons using AlphaSim (106). We identified clusters comprising more than 501 voxels as significant, with a global α level of 0.05.

### Data and code availability statements

The data and reconstruction toolbox presented in this manuscript will be available after we establish Material Transfer Agreement (MTA) between user’s institute and University of Southern California.

## 5. Acknowledgments

This work was supported by the National Institute of Health (NIH) grant UH3-NS100614, S10-OD025312, R01-NS114382, R01-EB028297, R21-EY028721, RF1-NS122028, R01-AG068055, P30-AG072946, R01-MH16948 and RF1-NS114628.

