## Supplemental information for "Age-Related Decline in Blood-Brain Barrier Function is More Pronounced in Males than Females in Parietal and Temporal Regions"

**Supplemental table S1**

| **Brain regions** | **Threshold age (years)** | **Age×Sex (F: 0, M: 1)**  (**Estimate** [95% CI]; **P value**) | **Slope for female**  (**Estimate** [95% CI]; **P value**) | **Slope for Male**  (**Estimate**) |
| --- | --- | --- | --- | --- |
| **Gray matter** | 62 | N.A. (N.S.) | **-0.82** [-1.35, -0.28]; **0.003** | **-0.82** |
| **White matter** | 62 | N.A. (N.S.) | **-0.87** [-1.42, -0.33]; **0.002** | **-0.87** |
| **Frontal lobe** | N.A. | N.A. (N.S.) | N.A. (N.S.) | N.A. |
| **Temporal lobe** | N.A. | **-0.22** [-0.34, -0.1]; **0.001** | N.A. (N.S.) | **-0.22** |
| **Parietal lobe** | 19 | **-0.2** [-0.35, -0.04]; **0.01** | **-0.33** [-0.76, -0.3]; **0.01** | **-0.53** |
| **ACC** | 36 | N.A. (N.S.) | **-0.35** [-0.59, -0.11]; **0.005** | **-0.35** |
| **PCC** | 19 | N.A. (N.S.) | **-0.41** [-0.63, -0.18]; **0.001** | **-0.41** |
| **Precuneus** | 19 | N.A. (N.S.) | **-0.36** [-0.57, -0.16]; **0.001** | **-0.36** |
| **Caudate** | 62 | N.A. (N.S.) | **-0.83** [-1.41, -0.24]; **0.006** | **-0.83** |
| **Putamen** | N.A. | N.A. (N.S.) | N.A. (N.S.) | N.A. |
| **Amygdala** | N.A. | N.A. (N.S.) | N.A. (N.S.) | N.A. |
| **Hippocampus** | 19 | **-0.14** [-0.26, -0.02]; **0.02** | **-0.12** [-0.43, -0.09]; **0.003** | **-0.26** |
| **PHG** | N.A. | **-0.15** [-0.26, -0.04]; **0.006** | N.A. (N.S.) | **-0.15** |
| **MTL** | N.A. | **-0.15** [-0.26, -0.04]; **0.008** | N.A. (N.S.) | **-0.15** |

**Supplemental Table S1: Analysis of age and sex dependent kw variations in 14 brain regions using MARS.** Threshold ages when kw starts declining have been identified. The age×sex interaction terms being negative suggest a more pronounced decline in kw in males compared to females. For females, the post-threshold age kw decline slope is presented with 95% confidence intervals (CI) and P values. The male kw decline slopes were estimated from the female slope and the age×sex interaction. Entries labeled 'N.A. (N.S.)' indicate non-significant changes.

**Supplemental table S2**

| **Brain regions** | **Threshold age (years)** | **Sex effect (F: 0, M: 1)**  (Estimate (95% CI); P value) | **Slope for female and male**  (**Estimate** [95% CI]; **P value**) | |
| --- | --- | --- | --- | --- |
|  |  |  | Before threshold age | After threshold age |
| **Gray matter** | 22 | **-8.91** [-11.94, -5.88]; **<0.001** | **-1.39** [-2.01, -0.77]; **<0.001** | **-0.39** [-0.47, -0.31]; **<0.001** |
| **White matter** | 22 | **-7.81** [-10.57, -5.06]; **<0.001** | **-1.35** [-1.92, -0.79]; **<0.001** | **-0.27** [-0.34, -0.2]; **<0.001** |
| **Frontal lobe** | 22 | **-8.43** [-11.60, -5.25]; **<0.001** | **-1.52** [-2.17, -0.87]; **<0.001** | **-0.41** [-0.5, -0.33]; **<0.001** |
| **Temporal lobe** | 22 | **-9.36** [-12.56, -6.16]; **<0.001** | **-1.4** [-2.06, -0.75]; **<0.001** | **-0.41** [-0.49, -0.32]; **<0.001** |
| **Parietal lobe** | 22 | **-10.77** [-14.11, -7.43]; **<0.001** | **-1.56** [-2.25, -0.88]; **<0.001** | **-0.46** [-0.55, -0.37]; **<0.001** |
| **ACC** | 22 | **-8.01** [-11.44, -4.59]; **<0.001** | **-1.19** [-1.89, -0.48]; **0.001** | **-0.46** [-0.55, -0.36]; **<0.001** |
| **PCC** | 22 | **-10.97** [-15.16, -6.78]; **<0.001** | **-1.83** [-2.69, -0.97]; **<0.001** | **-0.46** [-0.58, -0.35]; **<0.001** |
| **Precuneus** | 22 | **-10.65** [-14.14, -7.16]; **<0.001** | **-1.77** [-2.49, -1.05]; **<0.001** | **-0.39** [-0.49, -0.3]; **<0.001** |
| **Caudate** | 29 | **-5.28** [-7.52, -3.04]; **<0.001** | **-0.82** [-1.09, -0.54]; **<0.001** | **-0.23** [-0.3, -0.16]; **<0.001** |
| **Putamen** | 22 | **-4.75** [-7.32, -2.17]; **<0.001** | **-1.01** [-1.54, -0.48]; **<0.001** | **-0.11** [-0.18, -0.04]; **0.001** |
| **Amygdala** | 52 | **-5.35** [-8.16, -2.53]; **<0.001** | **-0.15** [-0.29, -0.02]; **0.03** | **-0.33** [-0.51, -0.15]; **<0.001** |
| **PHG** | 52 | **-5.75** [-8.45, -3.04]; **<0.001** | **-0.13** [-0.26, 0]; **0.06** | **-0.43** [-0.6, -0.26]; **<0.001** |
| **MTL** | 52 | **-5.59** [-8.17, -3.02]; **<0.001** | **-0.17** [-0.29, -0.04]; **0.008** | **-0.37** [-0.53, -0.2]; **<0.001** |
| **Brain regions** | **Threshold age (years)** | **Age×Sex (F: 0, M: 1)**  (Estimate (95% CI); P value) | **Slope for female and male**  (**Estimate** [95% CI]; **P value**) | |
|  |  |  | Before threshold age | After threshold age |
| **Hippocampus** | 22 | **-0.1** [-0.14, -0.05]; **<0.001** | **F**: **-0.74** [-1.35, -0.32]; **0.002**  **M**: **-0.84** | **F**: **-0.17** [-0.34, -0.19]; **<0.001**  **M**: **-0.27** |

**Supplemental Table S2: Analysis of age and sex dependent CBF variations in 14 brain regions using MARS.** Threshold ages where CBF slope changes occur were identified. A negative age×sex interaction term was detected only in the hippocampus, suggesting a more pronounced CBF decline in males compared to females. In other brain regions, the rate of CBF decline with age is largely similar between males and females, although males consistently exhibit lower CBF. For both males and females, the CBF decline slopes before and after the threshold age are presented with 95% confidence intervals (CIs) and P values.

**Supplemental table S3**

| **Brain regions** | **Threshold age (years)** | **Sex effect (F: 0, M: 1)**  (Estimate (95% CI); P value) | **Slope for female and male**  (**Estimate** [95% CI]; **P value**) | |
| --- | --- | --- | --- | --- |
|  |  |  | Before threshold age | After threshold age |
| **Gray matter** | 36 | **83.15** [37.19, 129.12]; **<0.001** | **N.A. (N.S.)** | **5.55** [3.83, 7.27]; **<0.001** |
| **White matter** | 36 | **73.93** [28.18, 119.68]; **0.002** | **N.A. (N.S.)** | **5.1** [3.39, 6.8]; **<0.001** |
| **Frontal lobe** | 36 | **84.99** [38.66, 131.32]; **<0.001** | **N.A. (N.S.)** | **5.67** [3.94, 7.4]; **<0.001** |
| **Temporal lobe** | 69 | **91.12** [42.21, 140.03]; **<0.001** | **5.49** [4.26, 6.72]; **<0.001** | **N.A. (N.S.)** |
| **Parietal lobe** | 43 | **N.A. (N.S.)** | **N.A. (N.S.)** | **6.9** [5.06, 8.74]; **<0.001** |
| **ACC** | 36 | **95.01** [43.55, 146.46]; **<0.001** | **N.A. (N.S.)** | **6.43** [4.89, 7.97]; **<0.001** |
| **PCC** | 36 | **N.A. (N.S.)** | **N.A. (N.S.)** | **5.55** [3.84, 7.25]; **<0.001** |
| **Precuneus** | 36 | **N.A. (N.S.)** | **N.A. (N.S.)** | **6.08** [4.53, 7.63]; **<0.001** |
| **Caudate** | 36 | **N.A. (N.S.)** | **N.A. (N.S.)** | **4.88** [3.48, 6.27]; **<0.001** |
| **Putamen** | 36 | **80.03** [31.87, 128.18]; **0.001** | **N.A. (N.S.)** | **5.27** [3.83, 6.71]; **<0.001** |
| **Amygdala** | 36 | **61.81** [16.47, 107.16]; **0.008** | **0.52** [-3.31, 4.34]; **0.79** | **5.0** [3.3, 6.69]; **<0.001** |
| **Hippocampus** | 52 | **N.A. (N.S.)** | **N.A. (N.S.)** | **6.5** [4.29, 8.71]; **<0.001** |
| **PHG** | 36 | **N.A. (N.S.)** | **N.A. (N.S.)** | **4.04** [2.78, 5.3]; **<0.001** |
| **MTL** | 52 | **N.A. (N.S.)** | **N.A. (N.S.)** | **6.45** [4.33, 8.57]; **<0.001** |

**Supplemental Table S3: Analysis of age and sex dependent ATT variations in 14 brain regions using MARS.** Threshold ages where ATT slope changes occur were identified. The rate of ATT increase with age is largely similar between males and females, although males consistently exhibit longer ATT. For both males and females, the ATT increase slopes before and after the threshold age are presented with 95% confidence intervals (CIs) and P values.

**Supplemental table S4**

| **Brain regions** | **Location**  **(mm)** | **Cluster size (voxels)** | **T value** | **P value** |
| --- | --- | --- | --- | --- |
| **Prefrontal cortex** | (120, 122, 124) | 9026 | -2.55 | 0.022 |
| **Cingulate cortex** | (90, 160, 66) | 2543 | -2.47 | 0.024 |
| **Precuneus** | (92, 64, 118) | 2938 | -2.42 | 0.024 |
| **Lateral temporal lobe** | (108, 48, 82) | 8598 | -2.60 | 0.020 |
| **Occipital lobe** | (110, 48, 86) | 3967 | -2.60 | 0.020 |
| **Insula** | (48, 128. 74) | 1878 | -2.75 | 0.017 |

**Supplemental Table S4: Voxel-wise analysis of age trend in kw.** Brain regions with significant negative correlations between kw and age with corresponding location (AAL template), cluster size, T values and P values.

**Supplemental table S5**

| **Measurement** | **Brain regions** | **Location**  **(mm)** | **Cluster size (voxels)** | **T value** | **P value** |
| --- | --- | --- | --- | --- | --- |
| **kw** | **Lateral prefrontal cortex** | (34, 118, 124) | 1055 | -2.22 | 0.031 |
|  | **Parietal lobe** | (128, 82, 122) | 1375 | -2.23 | 0.030 |
|  | **Lateral and medial temporal lobe** | (144, 66, 80) | 1517 | -2.24 | 0.030 |
| **CBF** | **Supra-marginal gyrus** | (144, 86, 100) | 584 | -2.27 | 0.029 |
|  | **Hippocampus** | (126, 110, 56) | 560 | -2.35 | 0.026 |
|  | **Frontal lobe** | (142, 142, 84) | 588 | -2.37 | 0.025 |
| **ATT** | **Supra marginal gyrus** | (122, 74, 122) | 1967 | 2.30 | 0.027 |
|  | **Posterior temporal lobe** | (144, 70, 82) | 4016 | 2.54 | 0.019 |
|  | **Calcarine sulcus** | (110, 56, 82) | 1531 | 2.33 | 0.026 |

**Supplemental Table S5: Voxel-wise analysis of age and sex effect in kw, CBF and ATT.** Brain regions with significant age×sex effects in kw, CBF and ATT with corresponding locations (AAL template), cluster sizes, T values and P values.

**Supplemental table S6**

| **Brain regions** | **CBF**  **Estimate** [95% CI];  **P value** | | **ATT**  **Estimate** [95% CI];  **P value** | |
| --- | --- | --- | --- | --- |
|  | **8 - 61 years** | **62 - 92 years** | **8 - 61 years** | **62 - 92 years** |
| **Gray matter** | **0.03** [-0.30, 0.35]; **0.88** | **0.43** [-0.15, 1.01]; **0.14** | **-0.03** [-0.05 -0.01]; **0.01** | **-0.07** [-0.11, -0.03]; **<0.001** |
| **White matter** | **-0.01** [-0.39, 0.37]; **0.97** | **0.31** [-0.32, 0.93]; **0.33** | **-0.02** [-0.05, 0.00]; **0.08** | **-0.07** [-0.11, -0.03]; **<0.001** |
| **Frontal lobe** | **-0.03** [-0.35, 0.30]; **0.87** | **0.18** [-0.42, 0.77]; **0.56** | **-0.02** [-0.05, 0.00]; **0.08** | **-0.07** [-0.11, -0.03]; **<0.001** |
| **Temporal lobe** | **0.09** [-0.23, 0.40]; **0.58** | **0.51** [-0.03, 1.04]; **0.06** | **-0.03** [-0.05, -0.01]; **0.01** | **-0.05** [-0.08, -0.01]; **<0.01** |
| **Parietal lobe** | **0.15** [-0.23, 0.53]; **0.44** | **0.40** [-0.23, 1.03]; **0.21** | **-0.03** [-0.05, 0.00]; **0.053** | **-0.03** [-0.08, 0.02]; **0.20** |
| **ACC** | **-0.20** [-0.56, 0.15]; **0.26** | **0.21** [-0.44, 0.85]; **0.52** | **-0.01** [-0.04, 0.01]; **0.33** | **-0.05** [-0.09, -0.01]; **0.007** |
| **PCC** | **0.17** [-0.19, 0.54]; **0.35** | **0.46** [-0.07, 0.98]; **0.09** | **-0.03** [-0.06, 0.00]; **0.10** | **-0.03** [-0.07, 0.01]; **0.09** |
| **Precuneus** | **0.03** [-0.37, 0.42]; **0.90** | **0.32** [-0.29, 0.92]; **0.30** | **-0.03** [-0.06, -0.00]; **0.03** | **-0.03** [-0.07, 0.01]; **0.16** |
| **Caudate** | **-0.45** [-0.93, 0.03]; **0.06** | **0.45** [-0.40, 1.30]; **0.29** | **-0.02** [-0.04, 0.01]; **0.31** | **-0.05** [-0.09, -0.01]; **0.007** |
| **Putamen** | **-0.54** [-1.01, -0.08]; **0.02** | **0.17** [-0.49, 0.82]; **0.61** | **0.00** [-0.03, 0.02]; **0.86** | **-0.05** [-0.08, -0.01]; **0.006** |
| **Amygdala** | **-0.57** [-0.98, -0.17]; **0.006** | **0.06** [-0.60, 0.71]; **0.87** | **0.01** [-0.02, 0.03]; **0.64** | **-0.06** [-0.09, -0.02]; **0.002** |
| **Hippocampus** | **-0.59** [-0.97, -0.22]; **0.002** | **0.15** [-0.55, 0.85]; **0.67** | **0.00** [-0.03, 0.02]; **0.80** | **-0.07** [-0.10, -0.03]; **<0.001** |
| **PHG** | **-0.50** [-0.85, -0.16]; **0.004** | **-0.03** [-0.66, 0.61]; **0.94** | **-0.01** [-0.03, 0.02]; **0.50** | **-0.07** [-0.11, -0.03]; **<0.001** |
| **MTL** | **-0.54** [-0.90, -0.18]; **0.003** | **0.09** [-0.59 0.77]; **0.79** | **-0.00** [-0.03, 0.02]; **0.65** | **-0.07** [-0.11, -0.04]; **<0.001** |

**Supplemental Table S6: Association between kw and CBF or ATT in 14 brain regions for participants aged between 8 - 61 years and 62 - 92 years.** Linear regressions incorporating sex as a covariate were conducted, and the estimated coefficients are presented with 95% confidence intervals (CI) and P values.

**Supplemental Figure S1**

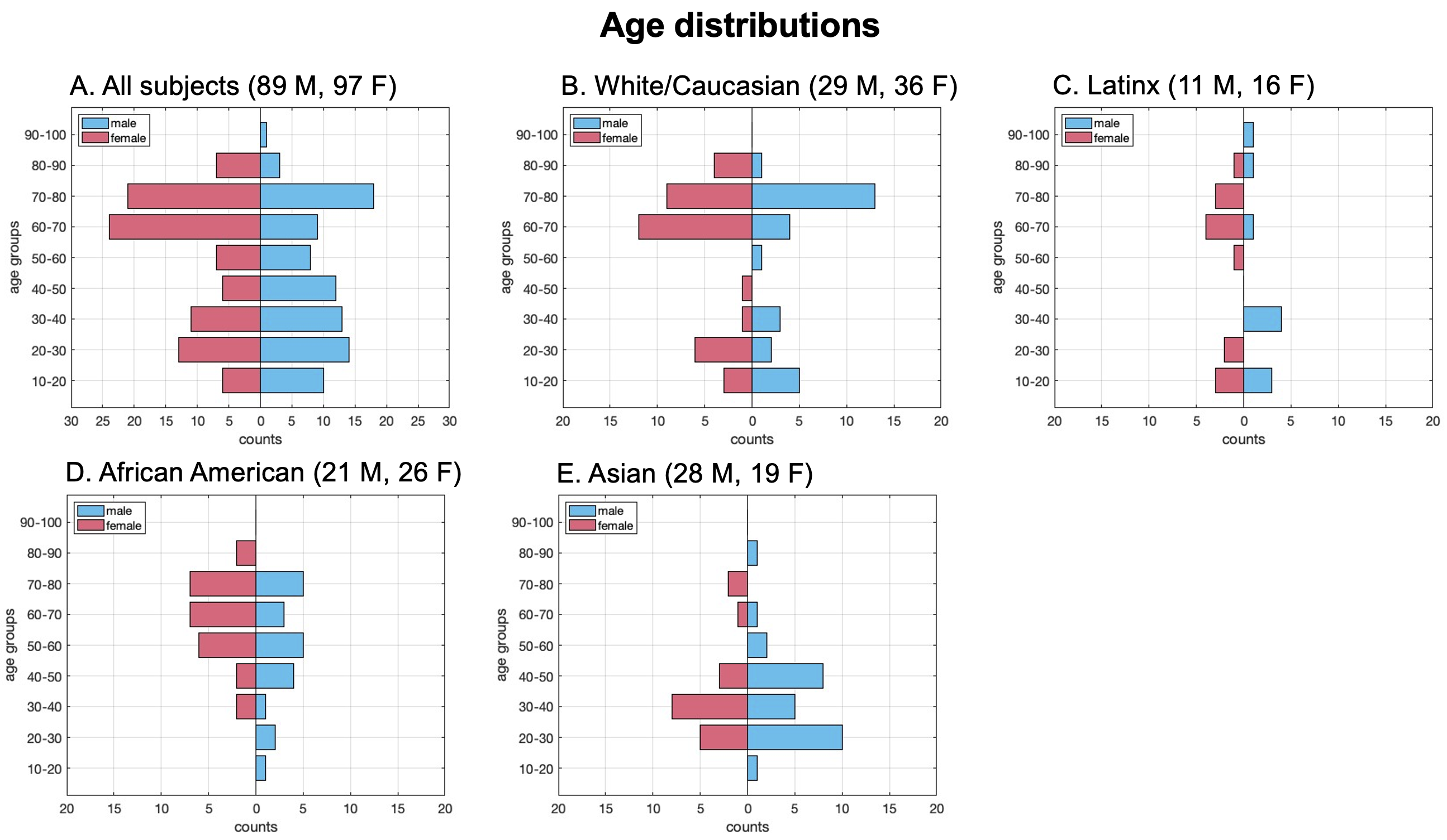

**Supplemental Figure S1.** Participant age distribution by sex. The study included 186 participants aged 8 to 92 years (89 males and 97 females). The racial distribution was as follows: 65 White/Caucasian (29 males, 36 females), 27 Latinx (11 males, 16 females), 47 African American (21 males, 26 females), and 47 Asian (28 males, 19 females). Since race may influence the changes in CBF, ATT, and kw with aging, it was incorporated as a covariate in the statistical analysis.

**Supplemental Figure S2**

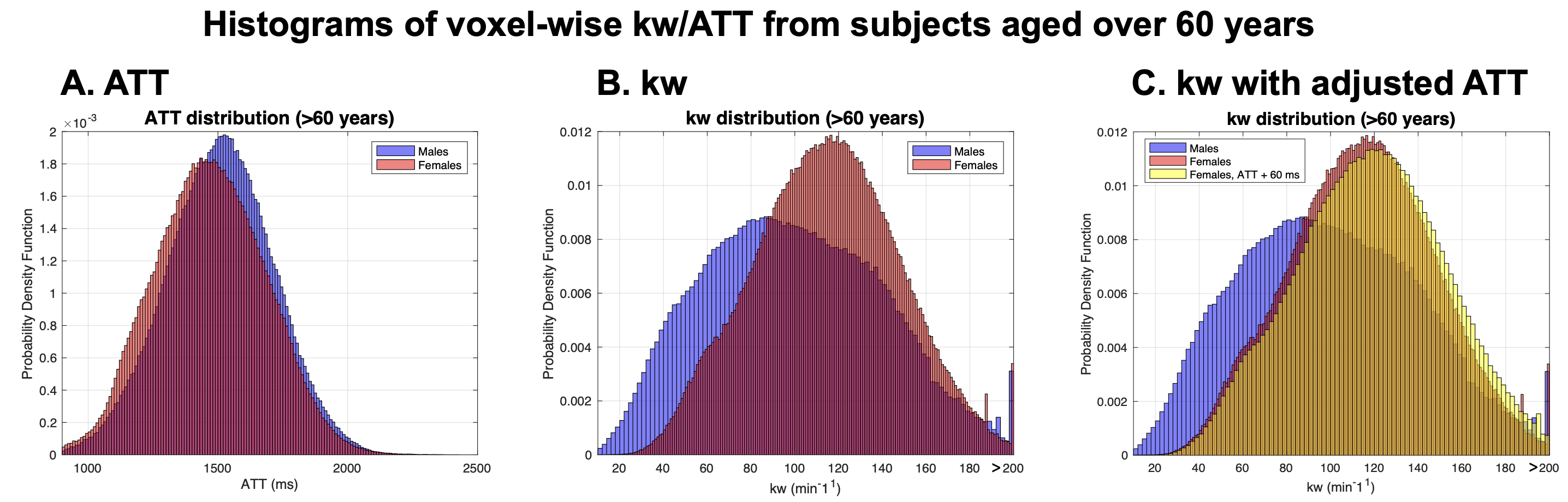

**Supplemental Figure S2.** Voxel-wise distributions of ATT and kw for participants aged over 62 years. (A) ATT distribution demonstrates a slightly longer ATT values in males (blue) than females (red), with similar distribution widths. (B) kw distribution reveals a higher mean kw in females (red), while males (blue) showing greater variability. (C) Simulated kw distribution for females with ATT adjusted by +60 ms, aligning with male ATT, indicates a marginally higher kw values compared to. These patterns suggest that differences in kw between males and females are not directly attributable to ATT variations. CBF values were not included in the simulations, as they do not affect kw quantification in SPA model.

**Supplemental Figure S3**

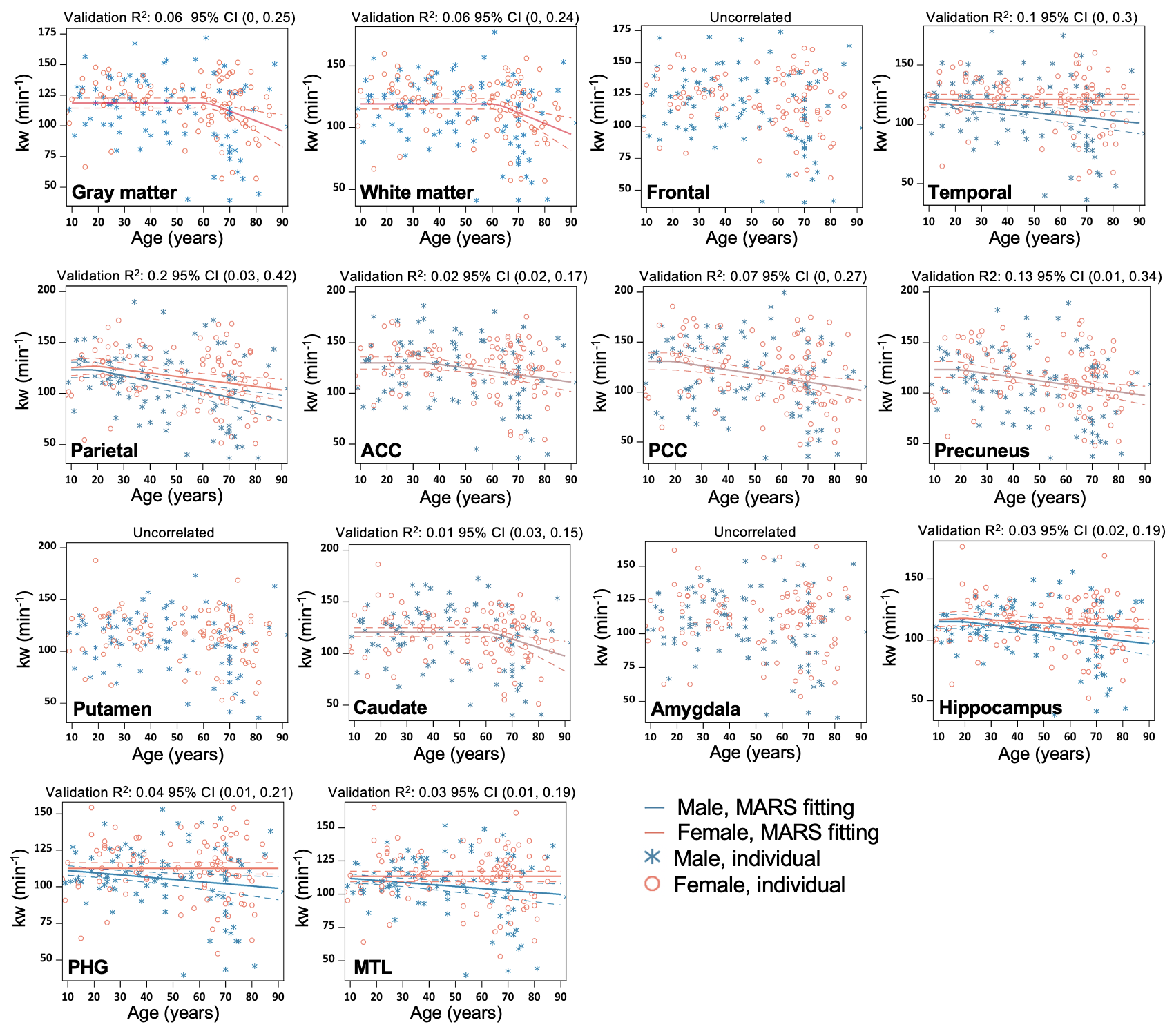

**Supplemental Figure S3.** Regional MARS analysis to reveal the age and sex dependent trajectories of kw in GM, WM, Frontal lobe, Temporal lobe, Parietal lobe, Anterior Cingulate Cortex (ACC), Posterior Cingulate Cortex (PCC), Precuneus, Putamen, Caudate, Amygdala, Hippocampus, Parahippocampal gyrus (PHG) and Mediotemporal lobe (MTL). Individual data points for males and females are indicated by blue asterisk symbols and red circles, respectively, while the corresponding MARS fitting curves (solid lines) and 95% confidence interval for predicted value (dash lines) are presented in matching colors. Validation R^2^ indicates the correlation between observed value and predicted value obtained through cross-validation.

**Supplemental Figure S4**

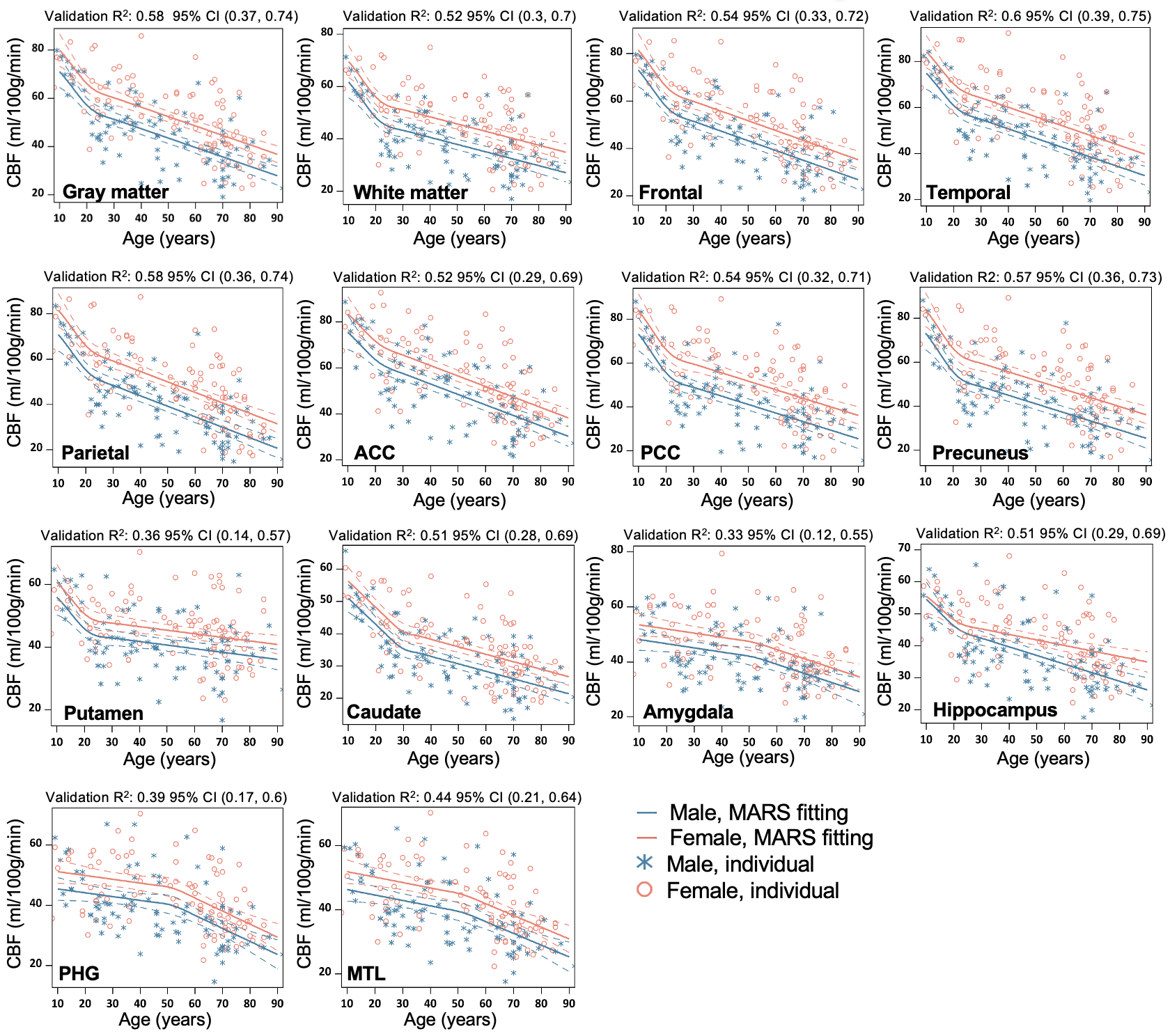

**Supplemental Figure S4.** Regional MARS analysis to reveal the age and sex dependent trajectories of CBF in GM, WM, Frontal lobe, Temporal lobe, Parietal lobe, Anterior Cingulate Cortex (ACC), Posterior Cingulate Cortex (PCC), Precuneus, Putamen, Caudate, Amygdala, Hippocampus, Parahippocampal gyrus (PHG) and Mediotemporal lobe (MTL). Individual data points for males and females are indicated by blue asterisk symbols and red circles, respectively, while the corresponding MARS fitting curves (solid lines) and 95% confidence interval for predicted value (dash lines) are presented in matching colors. Validation R^2^ indicates the correlation between observed value and predicted value obtained through cross-validation.

**Supplemental Figure S5**

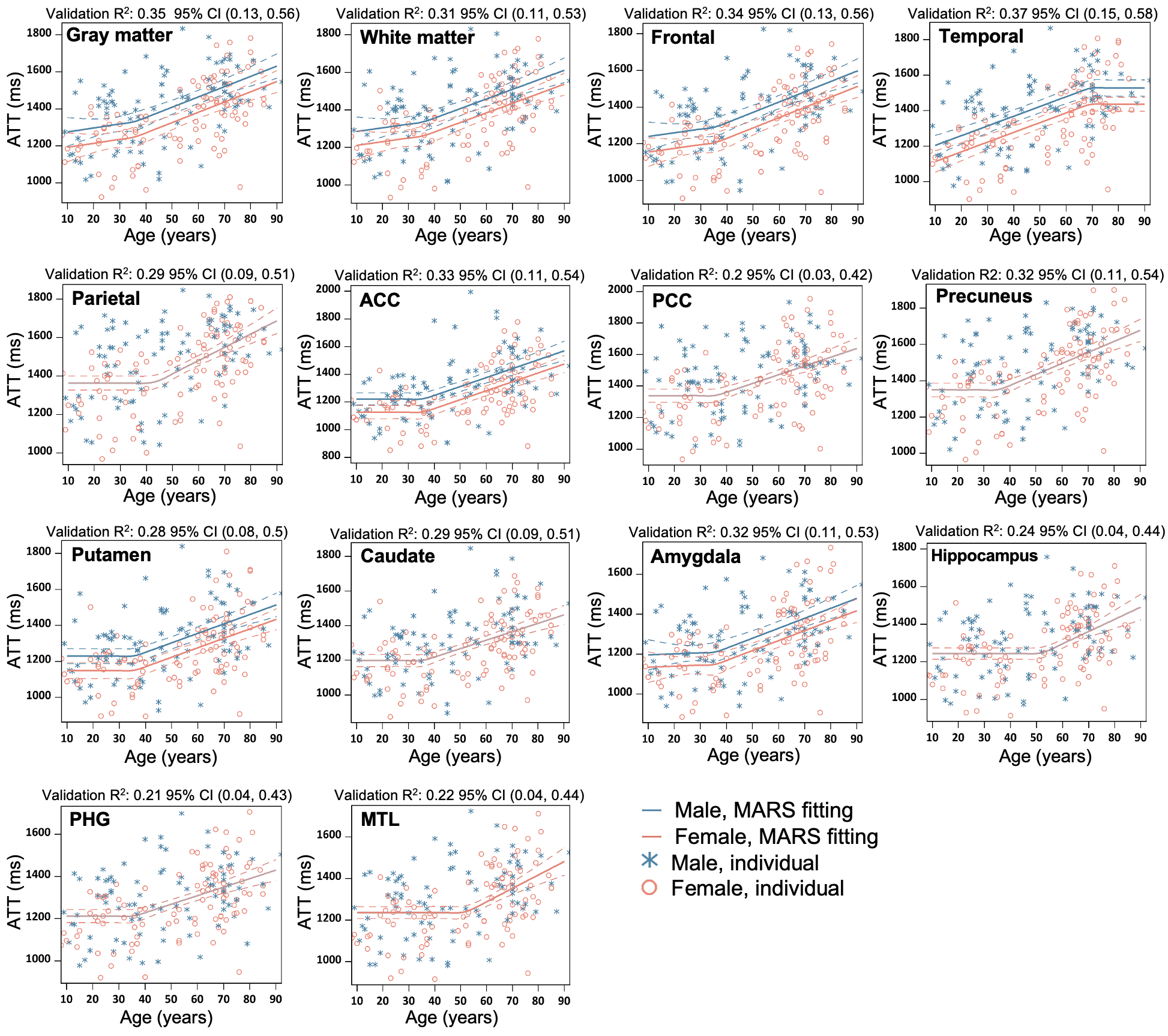

**Supplemental Figure S5.** Regional MARS analysis to reveal the age and sex dependent trajectories of ATT in GM, WM, Frontal lobe, Temporal lobe, Parietal lobe, Anterior Cingulate Cortex (ACC), Posterior Cingulate Cortex (PCC), Precuneus, Putamen, Caudate, Amygdala, Hippocampus, Parahippocampal gyrus (PHG) and Mediotemporal lobe (MTL). Individual data points for males and females are indicated by blue asterisk symbols and red circles, respectively, while the corresponding MARS fitting curves (solid lines) and 95% confidence interval for predicted value (dash lines) are presented in matching colors. Validation R^2^ indicates the correlation between observed value and predicted value obtained through cross-validation.
